# Network medicine informed multi-omics integration identifies drug targets and repurposable medicines for Amyotrophic Lateral Sclerosis

**DOI:** 10.1101/2024.03.27.586949

**Authors:** Mucen Yu, Jielin Xu, Ranjan Dutta, Bruce Trapp, Andrew A. Pieper, Feixiong Cheng

## Abstract

Amyotrophic Lateral Sclerosis (ALS) is a devastating, immensely complex neurodegenerative disease by lack of effective treatments. To date, the challenge to establishing effective treatment for ALS remains formidable, partly due to inadequate translation of existing human genetic findings into actionable ALS-specific pathobiology for subsequent therapeutic development. This study evaluates the feasibility of network medicine methodology via integrating human brain-specific multi-omics data to prioritize drug targets and repurposable treatments for ALS. Using human brain-specific genome-wide quantitative trait loci (x-QTLs) under a network-based deep learning framework, we identified 105 putative ALS-associated genes enriched in various known ALS pathobiological pathways, including regulation of T cell activation, monocyte differentiation, and lymphocyte proliferation. Specifically, we leveraged non-coding ALS loci effects from genome-wide associated studies (GWAS) on brain-specific expression quantitative trait loci (QTL) (eQTL), protein QTLs (pQTL), splicing QTL (sQTL), methylation QTL (meQTL), and histone acetylation QTL (haQTL). Applying network proximity analysis of predicted ALS-associated gene-coding targets and existing drug-target networks under the human protein-protein interactome (PPI) model, we identified a set of potential repurposable drugs (including Diazoxide, Gefitinib, Paliperidone, and Dimethyltryptamine) for ALS. Subsequent validation established preclinical and clinical evidence for top-prioritized repurposable drugs. In summary, we presented a network-based multi-omics framework to identify potential drug targets and repurposable treatments for ALS and other neurodegenerative disease if broadly applied.

## Introduction

Amyotrophic Lateral Sclerosis (ALS) is a neurodegenerative disease characterized by the death of motor neurons in spinal cords and brain resulting in skeletal muscle atrophy and eventually paralysis^1^. ALS with a prevalence rate of 5.2 per 100,000 individuals and affecting approximately 200,000 Americans, remains a highly sought-after topic in the forefront of both translational and clinical research^2^. Several potential pathological causes of ALS were reported, such as TDP-43 protein aggregation disrupting cell functions and leading to neuron death, glutamate excitotoxicity causing motor neuron degeneration and neuroinflammation^3–5^.

The underlying genetic factors of ALS have been explored by several national and international human genome sequencing projects^6^. For example, the ALS Consortium of the New York Genome Center and Answer ALS have generated multi-omics profiles, such as whole-genome sequencing, ATAC-sequencing, and RNA transcriptomics^6^. Furthermore, genome-wide association studies (GWAS) have identified dozens of genome-wide significant loci associated with risk of ALS^7^. With Edaravone and Riluzole being the only FDA-approved drugs, the task to translate available genetic and multi-omics data into efficacious therapeutics remains formidable, largely due to the complex, heterogenous pathobiology of ALS^8–10^. Oral masitinib has also shown clinical efficacy on both the primary and the secondary endpoints during the trial^11^. However, previous attempts of ALS therapeutics developments have not fully utilized or integrated the potential of genomics, protein-protein interactions, metabolomics, and transcriptomics - techniques that constitute the fundamentals of the emerging, interdisciplinary field of network medicine.^12,13^

The network medicine approach of understanding ALS involves the recognition that complex diseases, can be characterized by a multitude of genetic and environmental factors that interact within certain experimentally determined biological networks^12,13^. This approach effectively identifies molecular drivers in the human interactome and disturbed the cellular pathways that are involved in the pathogenesis and disease progression^14–17^. We therefore posit that unique integration of human genetics and multi-omics findings will offer new strategies to identify potential drug targets and repurposable treatments for ALS. Following these lines, we adopted a comprehensive, up-to-date, and multi-omics-dependent network topology-based deep learning framework^13^ to predict ALS-associated genes from large ALS GWAS findings (see Methods). The fundamental premise of our multi-omics framework was that likely risk genes of ALS exhibit distinct functional characteristics compared to non-risk genes and, therefore, can be distinguished by their aggregated brain-specific genome-wide quantitative trait loci (x-QTLs) features under the human protein-protein interactome network. In addition, we also demonstrated that multi-omics framework-predicted genes offer potential drug targets for emerging identification of repurposable treatments for ALS as validated by various preclinical and clinical models.

## Results

### A network-based multi-omics framework to predict drug targets for ALS

We adopted a network-based deep learning network constructed from integration of GWAS loci and human brain-specific functional genomics profiles (ref). Specifically, we leveraged non-coding ALS loci effects from GWAS on human brain-specific expression QTL (eQTL), protein QTLs (pQTL), splicing QTL (sQTL), methylation QTL (meQTL), and histone acetylation QTL (haQTL) under the human protein-protein interactome network model (*cf.* Methods). Generation of the network operates on the fundamental hypothesis that ALS-associated genes display unique functional attributes distinct from non-risk genes based on aggregated genomic features within the human protein interactome. As mentioned, we examined GWAS non-coding loci impacts on human brain-specific eQTL, pQTL, sQTL, meQTL, and haQTL (**Figure 1**). The execution of the network-based multi-omics framework are divided as follows: (1) sophisticated unsupervised deep learning model to partition protein-protein interactions (PPIs) into distinct functional network modules, capturing the inherent topological structures within the human protein interactome; (2) characterize each of these functional network modules by associating its nodes (proteins) with protein annotations derived from the Gene Ontology (GO) database^18^; (3) assess the significance of each node (gene) by integrating its functional similarity with genes identified through multiple brain-specific gene regulatory evidence (eQTL, pQTL, sQTL, meQTL, and haQTL), which have an influence on GWAS loci; (4) prioritize potential risk genes for ALS based on their collective gene regulatory features; and (5) predict repurposable drugs as potential therapy for ALS by network proximity analysis of multi-omics predicted ALS-associated gene-coding targets and existing drug-target networks under the human protein-protein interactome network (**Figure 1**).

**Figure 1.**
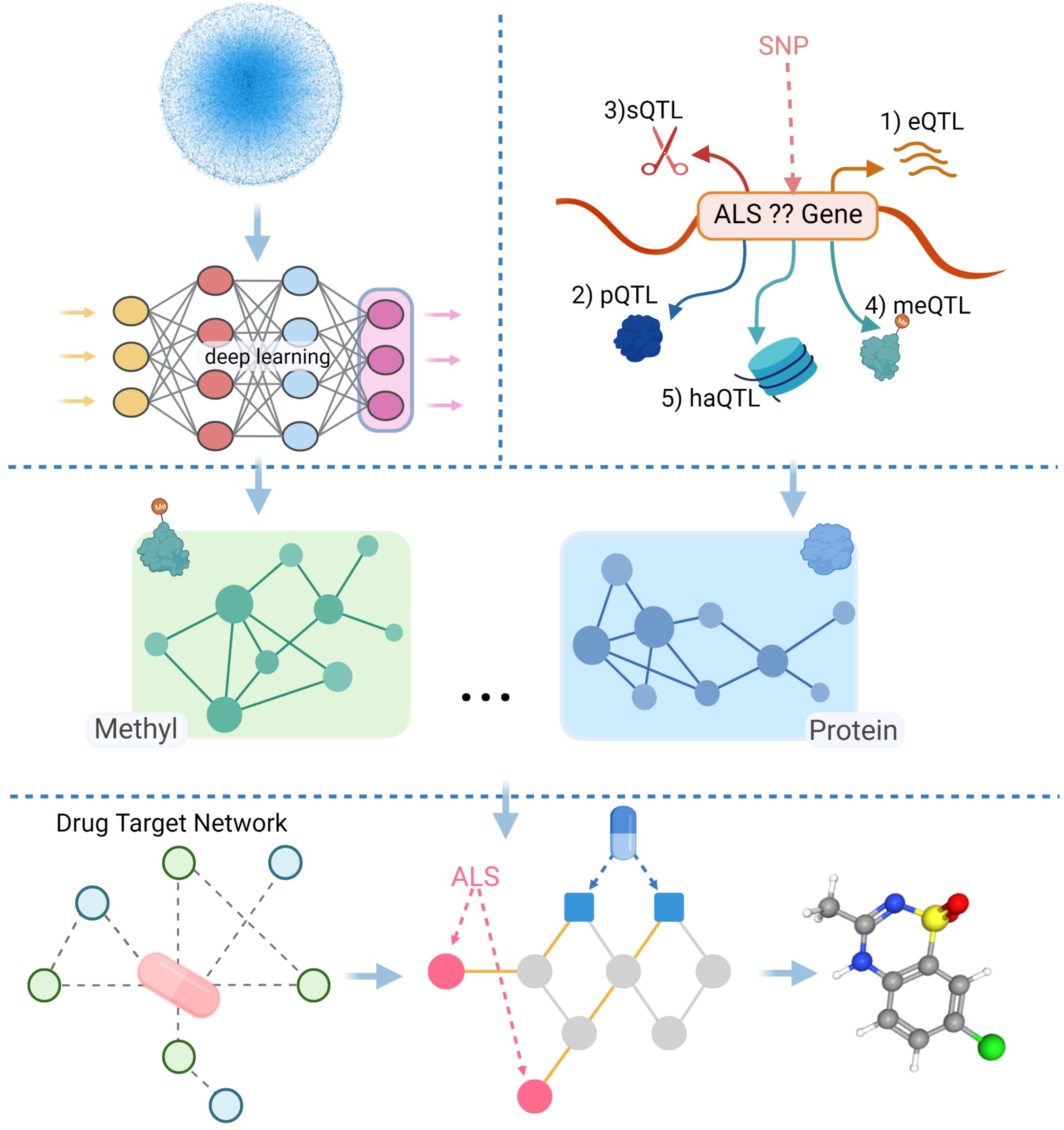
A flowchart describing the network-based multi-omics interaction workflow to infer drug targets and repurposable treatments for Amyotrophic Lateral Sclerosis (ALS). First, we employed an advanced machine learning technique to analyze the intricate network formed by protein-protein interactions (PPIs). This network was segmented into several smaller, interconnected clusters. We then found that these clusters could serve as predictors for protein roles as per the annotations in the Gene Ontology (GO) database. Moving forward, we identified potential genes associated with Amyotrophic Lateral Sclerosis. These genes share functional characteristics with previously known genes regulated by various genomic elements, such as methyl quantitative trait loci (meQTLs) and protein quantitative trait loci (pQTLs). In the final step, we focused on repurposing certain drugs (for example, gefitinib) that may be effective in treating Amyotrophic Lateral Sclerosis.

### A genome regulatory map of ALS GWAS loci

We examined the gene regulation involved in various ALS GWAS loci with human brain-specific eQTL, pQTL, sQTL, meQTL, and haQTL (**Figure 2**). Our analysis identified three genes with pQTL (*DHRS11, SARM1,* and *SCFD1*), 17 genes with haQTL (e.g., *SCFD1, MOBP, ERGIC1*), 4 genes with sQTL (*BAG6, CAMLG, TMEM175, FNBP1*), 109 genes with meQTL (e.g., *C9orf72, PTPRN2, SCFD1*), 93 genes with eQTL (e.g., *ATXN3, NEK1, C9orf72*). Notably, we identified several regulatory effectors targeting multitude of genes identified as ‘ALS loci’ candidates. For instance, Sec1 Family Domain Containing 1 (*SCFD1*), a significant risk factor for ALS patients, exhibits overlapping gene regulatory effect with eQTL, pQTL, haQTL, and meQTL^19^. BAG Cochaperone 6 (BAG6), which senses proteolytic fragments and prevents aggregation of TDP-43 fragments, exerts regulatory effect on eQTL, meQTL, and sQTL^20^. Myelin associated oligodendrocyte basic protein (MOBP), another prominent ALS risk gene, is enriched by eQTL, haQTL, and meQTL^21^. NIMA-related kinase 1 (*NEK1*), a susceptibility gene identified by GWAS, has gene regulatory influence on eQTL^22^. Overall, mapping of ALS loci into human brain-specific x-QTLs therefore showed the feasibility of integrating multi-omics data to identify the gene regulatory processes.

**Figure 2.**
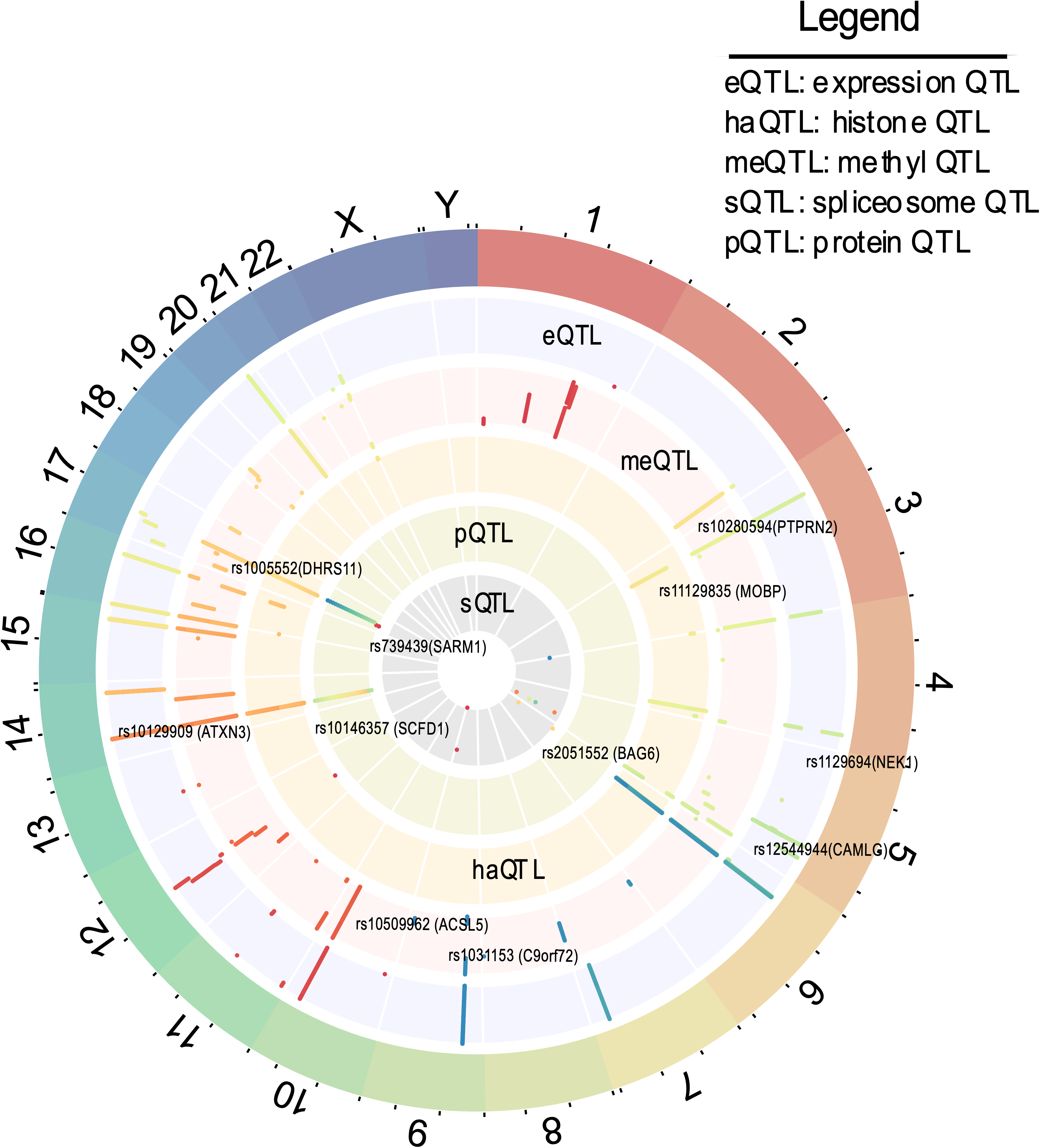
Gene regulatory landscape of ALS GWAS loci. Overview of genetic loci linked to Amyotrophic Lateral Sclerosis (ALS) identified through genome-wide association studies, distributed among various chromosomes and analyzed in relation to five genomic regulatory factors: expression quantitative trait loci (eQTL), histone modifications, protein interactions, spliceosome components, and DNA methylation patterns.

### Identification of likely ALS risk genes via multi-omics data integration

Clustering of PPIs into functional subnetwork modules using a topology-based self-learning framework (**Figure 1** and *cf.* Methods), identified groups of proteins that function together or are involved in similar biological processes. The resulting subnetwork modules provide a representation of biological relationships among proteins. Our previous work showed that proteins with more gene ontology (GO) terms tend to have a) more network clusters in the human PPI interactome^17^, and b) proteins with the same subnetwork module shared more GO annotation and a higher degree functional similarity. We integrated the PPI-derived network modules with five types of gene regulatory elements (brain-specific eQTL, pQTL, sQTL, meQTL, and haQTL) implicated by ALS GWAS Loci to determine to what extent which a specific gene is functionally related to the known ALS-associated loci. Taking haQTL as an example, we first assigned a predicted score to each gene based on its functional overlap with the histones. By evaluating the correlation between the predicted score and the presence of histones-linked genes associated with ALS, we are able assess the extent to which a specific gene is functionally related to the known ALS-associated loci. Subsequently, we generated a comprehensive prediction through integrating all five types of human brain-specific x-QTLs by summing their individual scores.

In total, we pinpointed 105 putative ALS-associated genes by applying a Z-score cutoff (*cf.* Methods) which included *SCFD1, EIF2AK3, NRG1, UBQLN1*, and *CHRNA4*, and others. Among the 105 ALS-associated gene-coding proteins, a total of 273 PPIs (termed ALS disease module) from the largest connected components in the human PPI (**Figure 3**) was identified. We next compared the constructed ALS PPI network with published ALS-associated genes from DiseNET^23^, Open Targets^24^, KEGG PATHWAY database, and other ALS patients’ brain transcriptomic data (see Methods). We found that the established ALS disease module is significantly enriched in the DisgeNET gene set (p = 0.0080, Fisher’s exact test), KEGG PATHWAY database (p = 0.043), as well as GWAS Catalog 2023 (p = 0.032). In addition, pathway enrichment analyses demonstrated that the 105 predicted ALS-associated genes were significantly enriched in a multitude of immune pathways, namely regulation of T cell activation (q = 1.07 x 10^-10^), monocyte differentiation (q = 1.15 x 10^-5^), and lymphocyte proliferation (q = 6.22 x 10^-6^). Overall, our results identified several ALS-related pathobiological pathways that are affected by the 105 ALS predicted genes.

**Figure 3.**
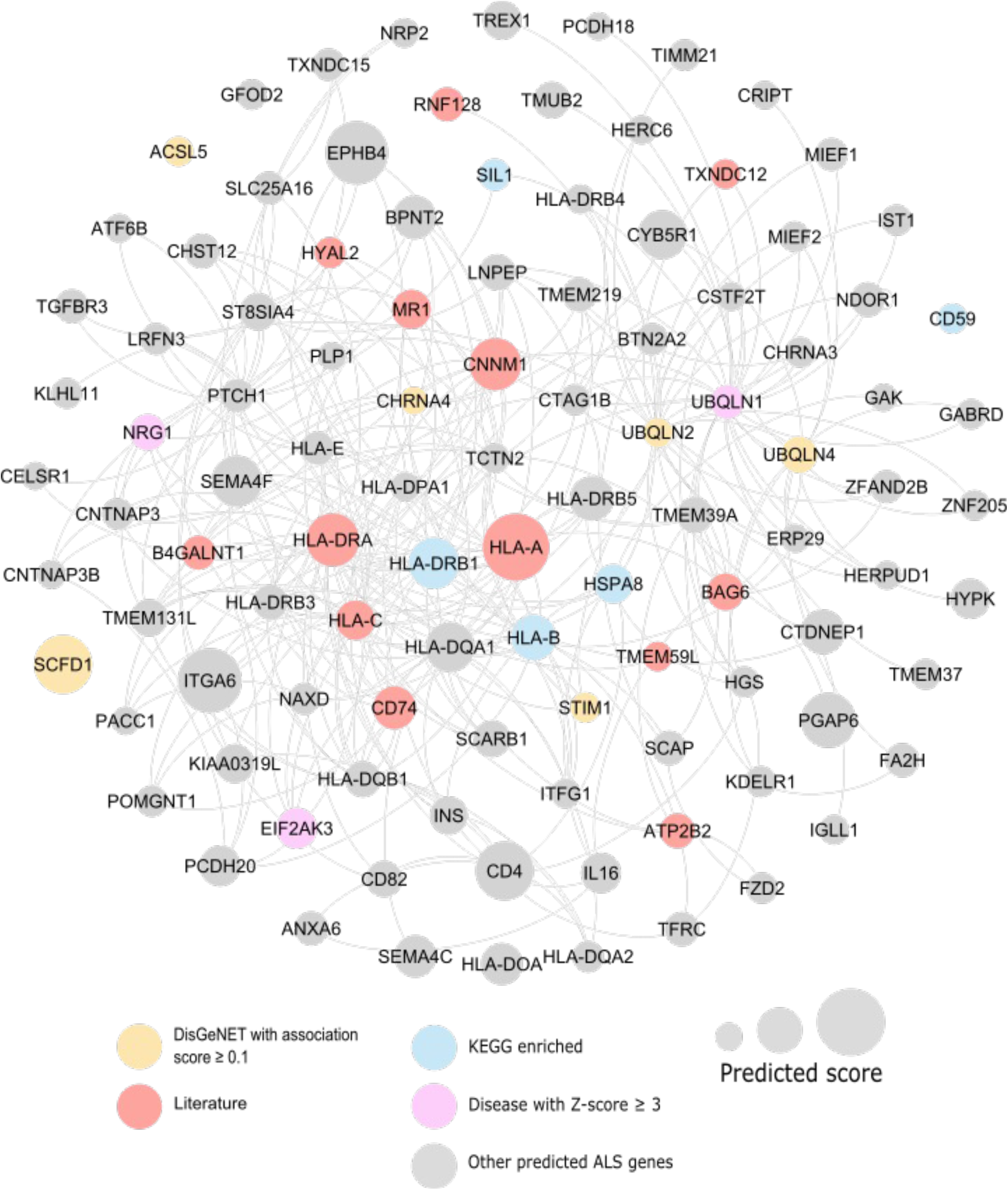
Network-based visualization of 105 predicted ALS-associated genes. Prioritized ALS-associated genes are colored with various evidence. Yellow genes are the ones identified by GisGeNET with an association score ≥ 0.1. Blue genes are the ones enriched in KEGG pathway. Purple geens are the ones detailed in Diseases JansenLab database with a Z-score ≥ 3. Red Genes are the ones identified by enrichment analysis from other ALS-relavant literature.

### Network-based discovery of potential repurposable drugs for ALS

We next assessed the efficacy of drugs predicted by our multi-omics framework and its applicability to other published ALS-associated gene databases. We assembled five additional ALS risk gene sets from several public sources, namely DisGeNET (Disgenet.org), Open Targets (opentarget.org), all ALS-connected genes from ALSoD, (alsod.ac.uk), Jensen Disease, and all ALS-associated genes from Refmap (see Methods). To screen for high confidence ALS-associated gene sets, we constructed an additional cumulative gene list of 155 genes, named “Union” (**Figure 5**), that includes genes that appeared at least twice across the five gene databases. Subsequently, we performed network proximity analysis for each of the five literature-supported ALS-associated gene lists as well as the Union gene list and compared their results with our multi-omics framework predicted ALS-associated genes. We found significant overlapping between our multi-omics predicted results and known ALS gene database-predicted drugs. Among those, Diazoxide is significantly enriched in all five individual literature gene lists and tops the Union gene list (Z < −2.0, p < 0.01). Dimethyltryptamine (DMT) is significantly enriched in Union (Z = −2.165, p = 0.012), DisGeNET (Z = −2.543, p = 0.009), Open Targets (Z = −2.032, p = 0.008), Refmap (Z = −2.173, p = 0.014). Gefitinib is significantly enriched in DisGeNET (Z = −2.43, p = 0.006), Jensen Disease (Z = −2.309, p = 0.016), Refmap (Z = −2.345, p = 0.007). Importantly, our network proximity analysis multi-omic data of predicted ALS disease modules also revealed novel findings (like Paliperidone) that were not identified in any of the five literature gene lists.

We next turned to perform the network proximity analyses of the predicted ALS-associated genes by calculating the closest distance between a drug’s binding targets and the ALS disease module on the human PPI network^12,16^. We prioritized candidate repurposing drugs using: (1) strong network proximity (Z-score < −2.0); (2) ideal brain penetration properties; (3) existing experimental evidence from functional studies or clinical trials. Using these criteria, we identified a total of 20 prominent candidates. Together, nine out of fourteen different classes (e.g., alimentary tract and metabolism, blood and blood forming organs, and nervous system) defined in the first level of the Anatomical Therapeutic Chemical (ATC) codes were represented across our 20 top prioritized candidate drugs. This highlights the potential of our multi-omics framework to expand the scope of ALS drug target discovery from large human genetics findings beyond existing knowledges.

Diazoxide (Z = −2.81) is a small FDA-approved drug known for activating ATP-sensitive potassium channels (KATP) present in neurons and glial cells within the Central Nervous System (CNS). Studies have shown Diazoxide to play a role in protecting dopaminergic neurons from death by reducing astrocyte and microglial activation and reducing neuroinflammation associated with activated microglia^25^. As in the context of neuroinflammation involving the release of pro-inflammatory mediators that leads to damage and dysfunction in the CNS, diazoxide may help regulate the inflammatory response and limit neurotoxicity due to its ability to cross the blood-brain barrier^26^. Our study illustrates the feasibility of above mechanism. For instance, a direct druggable target of diazoxide is *SLC12A3*, a family of transporters involved in reuptake of excess glutamate from synaptic cleft, and thus a likely inhibitor of the neurotoxic nature that is ALS^27^. Notably, *SLC12A3* is in primary association with HSPA8, one of our predicted ALS PPI nodes (**Figure 4b**). Recent clinical studies have further validated the potential of diazoxide in mitigating neuroinflammation and confirmed its primary role in neuroprotection mediated by maintenance of mitochondrial homeostasis, reduction of oxidative stress, and protection excitotoxicity^28^. Overall, diazoxide’s anti-inflammatory properties and neuroprotective makes it a potential drug candidate in treating neuroinflammation in ALS.

**Figure 4.**
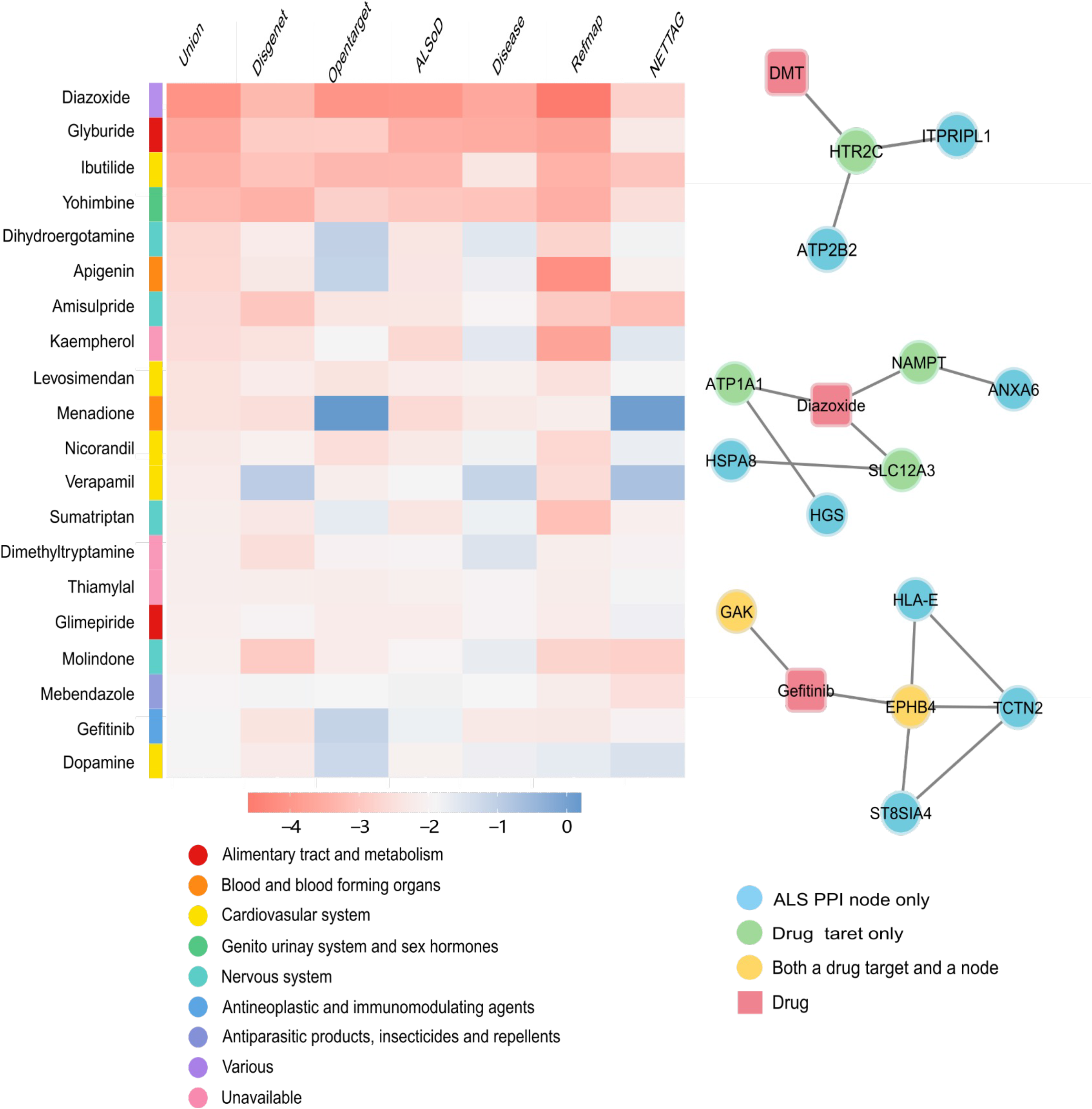
Network Proximity-predicted drugs for six existing gene sets from literatures and the predicted ALS-associated genes. Z-score between −4 to 0 is depicted by the gradient from red to blue. Drugs are categorized by colors according to the primary codes of the Anatomical Therapeutic Chemical (ATC) classification system. Three candidate drugs and their target genes using our drug-target network analysis. Blue genes are predicted ALS PPI nodes only, green genes are druggable targets that are in direct proximity with predicted ALS PPI nodes, yellow genes are both an ALS PPI node and a druggable target.

Gefitinib, primarily recognized for inhibiting epidermal growth factor receptor (EGFR) activity in lung cancer treatment, holds a prominent position (Z = −2.026) on the list. Notably, it demonstrates a positive impact on rescuing motor neuronal survival in a co-culture model of ALS, suggesting its potential neuroprotective effect by impeding the pathways responsible for progressive degeneration observed in ALS. Furthermore, gefitinib exhibits an enhanced reduction of TDP-43 proteinopathy, a significant hallmark of ALS disease onset, leading researchers to propose its modulation of the underlying cellular mechanisms involved in ALS. Moreover, gefitinib induces autophagic flux and facilitates cellular clearance^29^. Notably, GAK, or Cyclin G Associated Kinase, is both a direct drug target of gefitinib and one of our predicted ALS PPI nodes (**Figure 4c**). GAK is found to regulate autophagy through clearing misfolded and damaged organelles that are characteristic in affected neurons. Modulating GAK activity could potentially enhance the autophagic clearance of pathological in ALS, such as misfolded SOD1 or TDP-43^7^. Lastly, a different study has shown gefitinib to positively facilitate the PINK1/Parkin-mediated mitophagy, another hallmark of ALS pathobiology^30^. Combining gefitinib’s ability to pass through the blood-brain barrier, its neuroprotective influence on motor neurons, impact on astrocytes, TDP-43 regulation, and induction of autophagy, together they lend support to the hypothesis that gefitinib could play a therapeutic role in treating ALS^31^.

Paliperidone (Z = −2.153) is a drug that primarily treats schizophrenia through both the serotonin Type 2 (5HT2A) and the dopamine Type 2 (D2) receptor antagonism. It was recently found that paliperidone interacts with the amino acid residue Trp32 in the superoxide dismutase 1 (SOD1) protein, of which its mutation leads to unstable structures and aggregation, a major contributor to ALS pathogenesis. Specifically, paliperidone binds to the beta-strand 2 and 3 regions of the SOD1 protein, regions that play a role in scaffolding SOD1 fibrillation, the process by which SOD1 forms abnormal aggregates. By binding to these regions, paliperidone may disrupt the formation of SOD1 fibrils, potentially inhibiting aggregation^7^. Moreover, another study suggests that paliperidone in young-adult mice prenatally exposed to maternal immune challenge elicits a 411preventive anti-inflammatory and antioxidant effect in the frontal cortex. It is found to block the neuroinflammatory response and stimulates endogenous antioxidant mechanisms in the mice^32^.

Dimethyltryptamine (Z = −2.026) is a naturally occurring hallucinogenic compound worked primarily through activating the sigma-1 receptor, a protein that plays a role in survival and neuroplasticity. Dimethyltryptamine has been found to protect neurons from damage caused by glutamate toxicity, a key process that is through to play a role in the pathogenesis of ALS^33^. A different study found that DMT, through its interaction with the S1R receptor, can increase cellular survival by reducing hypoxic stress cultured in human cortical neurons, monocyte-derived macrophages, and dendritic cells^34^. Additionally, co-localization of the S1R receptor with the indolethylamine N-methyltransferase (INMT) in primate and human motor neurons (MNs) may provide protections in ALS. INMT is an enzyme involved in the biosynthesis of DMT. The presence of both S1R and INMT in MNs suggests that DMT may exert its protective effects in ALS through specific mechanisms that are yet to be discovered. Lastly, it was found that through enzyme activity of methylation of thiols and trace metals, DMT can reduce oxidative stress, a process known to contribute to the pathogenesis of ALS^35^.

## Discussion

In this study, we applied a network-based deep learning framework combined with large ALS GWAS loci and human brain-specific x-QTL findings to identify potential candidate drugs for ALS (**Figure 1**). Our study provides evidence that leveraging human brain-specific x-QTL data and combined with protein-protein interactome, is effective in inferring potential ALS drug targets and candidate repurposable treatments. Of note, we identified diazoxide, gefitinib, dimethyltryptamine, and paliperidone as potential ALS-repurposable therapeutics. Of critical importance is that, all four candidates can pass through the blood-brain barrier and have been shown to alleviate key ALS-pathobiological hallmarks such as neuroinflammation, supporting their potential efficacy in ALS treatment.

Recent human genomic technologies have witnessed several advances in the field of ALS-associated genetics, epidemiology, and therapy development that further enhances, and even reshapes, our current understanding about ALS pathobiology. One study focused on the characterization of missense variants in 24 ALS risk genes and found that these variants were enriched in a number of structural regions of proteins, such as beta-sheets and alpha-helices. These enrichments of variants in genes with high and medium expression level particularly within the brain, which provided insights into pathogenicity of missense variants in ALS and distinguish them from variants traditionally associated with neurodegenerative diseases^38^. Another study investigating the genetic overlap between Alzheimer’s disease (AD), Parkinson’s disease (PD), and ALS unveiled causal variants between ADRD and ALS at the TSPOAP1 locus, and between PD and ALS at the NEK1 and GAK/TREM175 loci^36^. Via comprehending the role of missense variants and shared underlying mechanisms, there is potential for significant implications in the development of targeted therapies and interventions that could benefit multiple neurodegenerative diseases simultaneously. This deeper understanding of the genetic basis and pathobiology of ALS enhances the prospects for the advancement of effective treatment strategies, offering hope for improved outcomes not only for ALS but also for related neurodegenerative disorders.

We acknowledged the limitation of this study being dependent on computational predictions and network analysis, which requires further experimental validation. While our multi-omics framework provides a comprehensive approach to identifying ALS risk genes and potential drug candidates, the true efficacy and safety of these drugs need to be further investigated in preclinical and clinical studies. Additionally, the study relies on the availability and accuracy of multi-omics data and protein-protein interaction networks. Any limitations or biases in these data sources could affect the reliability of the predictions. Further research is therefore needed to explore the therapeutic potential of the identified drugs and to understand their precise mechanisms of action in the context of ALS.

Despite these limitations, the study represents a significant step forward in the search for ALS treatments. By leveraging the power of network medicine and deep learning methodologies, it provides a systematic approach to identify ALS risk genes and repurposable drugs^12,13^. The identification of the four novel drug candidates highlights the potential of repurposing existing FDA-approved drugs for ALS treatment. Furthermore, the comparison with published findings validates the reliability of our multi-omics predictions and uncovers additional drug candidates that were not identified in previous studies. These findings open new avenues for therapeutic exploration and may contribute to the development of more effective treatments for ALS.

## Methods and Materials

### Description of a multi-omics framework

NETTAG included 3 steps. In the first step, we created a graph neural network (GNN)-based overlapping community detection methods to capture the PPI’s topology

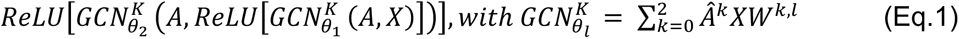

Here: 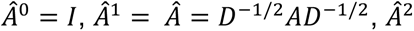 is the elementwise square of and the sub/super-script *l* is the layer index. A is the adjacency matrix of the PPI, D is the corresponding diagonal degree matrix, while X signifies the node feature matrix in general. Finally, the dimension of the final output layer interprets the clustering number. The output matrix is then feed into Bernoulli-Poisson model (Eq.2.1) to learn PPI’s topology (Eq.2.2). The output matrix has N (number of total nodes in PPI) rows and C (clustering numbers) columns. The final clustering number was determined as the one with lowest akaike information criterion (AIC) value (Eq.2.3). The loss explained itself as an overlapping-clustering method: if two nodes have multiple commonly shared clusters (is large), then there should exist an edge connecting each other ( is close to 1) and vice versa.

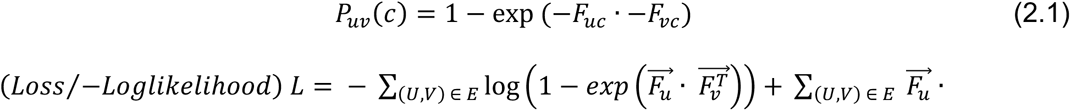

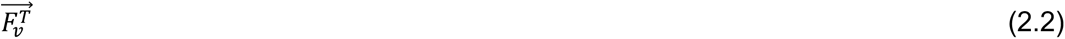

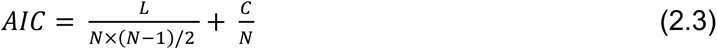

In the second step, we utilized clustering similarity to score each node with respect to each QTL. For example, assuming that we have 5 discovered ALS genes associated with eQTL, i.e. *SCFD1*, *CHRNA4*, *EIF2AK3*, *NRG1* and *UBQLN1*. Then for any given node, its score (Eq.3) based on eQTL is defined as:

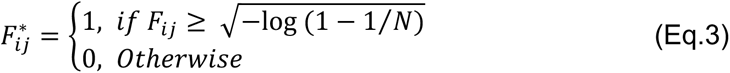

We repeated this procedure for all QTLs, therefore for each gene we have 5 different scores, and the final predicted score for each gene is the sum of all these 5 scores. Finally, in order to improve the robustness and stability of the clustering results, we run NETTAG with 10 different seeds and used ensembled prediction results as the final gene prioritization results (**Table S1**).

### Integration of human brain-specific functional genomic features

We assembled a total of 1,047,489 SNPs detailing various genetic components from the GWAS catalog^37^, including Amyotrophic Lateral Sclerosis. We then gathered regulatory elements data from five databases, namely Ensembl Regulatory Build, ENCODE, GTEx, and Roadmap and utilized the web-ready platform SNPnexus to annotate SNPs from the GRCh Build 38 human genome (GRCh38). In total, five gene regulatory elements (eQTL, pQTL, spliceosome, open chromatin, meQTL) were selected for assessment of ALS-associated genes (**Table S2**). Specifically, the process involves: first, we grouped together ALS SNPs detailed in GWAS Catalog with SNPs corresponding to each regulatory element, e.g., eQTL; second, the respective genes for each ALS-associated SNPs were pinpointed by the “MAPPED GENE(S)” column on GWAS catalog. Lastly, remaining SNPs with no matched ALS genes but are curated in other categorized genes (REPORTED GENE(S) column in GWAS) were mapped to its reported genes. For both Ensembl Regulatory Build and ENCODE, we only selected brain and normal karyotype and neuron tissues epigenetic components. For downstream analysis, we exclusively considered significant eSNPs (q < 0.05, LD r^2^ < 0.1) that were linked to eGenes when mapping eQTLs using the GWAS Catalog. These identified eSNPs were utilized as input features, although only one most significant eQTL feature were counted when multiple SNPs corresponding to the same eGene.

### Constructing the human protein-protein interactome

To construct the most up-to-date and comprehensive human PPI, we assembled data from commonly used bioinformatics databases consisting of five experimental assays based on experimental evidence: (i) binary PPIs predicted by high-throughput yeast-two-hybrid (Y2H) systems; (ii) physical, binary PPIs from tertiary protein structures from the Instruct knowledgebase^38^ (iii) kinase-substrate interactions evidenced by literature-derived high-throughput and low-throughput assays; (iv) signaling networks evidenced by literature-derived high-throughput and low-throughput experiments from the SignaLink2.0^39^. (v) literature-curated protein complex data predicted by affinity purification-mass spectrometry (AP-MS), or by other literature-derived low-throughput assays. Genes on the PPI network was mapped using their respective ENTREZ ID from the NCBI database. In total, our constructed human PPI network consists of 17,706 distinctive proteins with 351,444 PPIs. This study utilizes only the largest-connected components of our dataset, or 17,456 proteins and 336,549 PPIs.

### Literature-derived ALS-associated genes

To better assess the efficacy and applicability of our NETTAG-predicted therapeutics in comparison to published prominent findings, we compiled five additional lists of ALS risk genes from reputable sources, each selected based on specific criteria (**Table S3**). These sources included DisGeNET (top 100 genes with the highest disease-gene association score), Open Targets (genes with an overall association score ≥ 0.10), all ALS-connected genes from ALSoD, JensenLab Diseases (genes with a Z-score ≥ 4.7), and all ALS-associated genes from Refmap^23,24,40–42^. To identify more significant results, we created a cumulative gene list called “Union”. This list comprises 155 genes that appeared in at least two of the five gene lists mentioned above (**Table S3**). By combining these gene lists, we aimed to enhance the sensitivity and reliability of our analysis in identifying potential ALS risk genes.

### Network proximity analysis for drug prediction

In this study, FDA-approved and clinically investigated molecules were collected from the DrugBank database, which, combines with NETTAG-predicted ALS disease module, allowing us to perform the network proximity analysis with mechanisms below (**Table 4**).

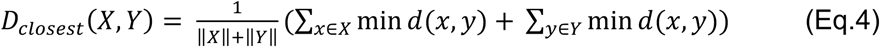

In our study, we used the notation d(x, y) to represent the shortest path length between protein x and protein y, belonging to the protein sets X and Y, respectively. Specifically, X represents the disease module obtained from NETTAG, while Y represents the set of drug targets associated with each drug, represented as proteins or genes. To assess the statistical significance of the proximity between these protein sets, we transformed the computed network proximity into a Z score using the following formula:

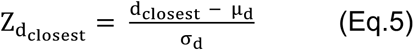

In our analysis, we utilized permutation tests with 1,000 random experiments to estimate the mean (μ_d_) and standard deviation (σ_d_). For each random experiment, we constructed two subnetworks, X_r_ and Y_r_, in the protein-protein interaction (PPI) network. These subnetworks were designed to have the same number of nodes and degree distribution as the original subnetworks, X and Y, respectively. The purpose of this approach was to mitigate any potential literature bias that may arise from well-studied proteins.

### Enrichment Analysis

We performed pathway and disease enrichment analyses using various databases. Specifically, we utilized GWAS Catalog 2023 and DisGeNET from the online platform Enrichr^43^ (**Table S5**). The combined score, as defined in Enrichr, was calculated by multiplying the logarithm of the p-value from the Fisher’s test with the Z score, which quantifies the deviation from the expected rank. The GWAS Catalog served as a valuable and easily accessible database of SNP-trait associations that have been identified through literature research. On the other hand, DisGeNET integrated information about genes associated with diseases. This comprehensive platform combines disease-associated genes from multiple sources, including expert-curated repositories, animal models, and scientific literature.

## Acknowledgements

**Funding** This work was primarily supported by the National Institute of Aging (NIA) under Award Number R21AG083003, R01AG084250, R56AG074001, U01AG073323, R01AG066707, R01AG076448, R01AG082118, and RF1AG082211, and the National Institute of Neurological Disorders and Stroke (NINDS) under Award Number RF1NS133812 to F.C. This work was supported in part by the Rebecca E. Barchas, MD, Professorship in Translational Psychiatry, the Valour Foundation, Project 19PABH134580006-AHA/Allen Initiative in Brain Health and Cognitive Impairment, the Elizabeth Ring Mather & William Gwinn Mather Fund, S. Livingston Samuel Mather Trust, and the Louis Stokes VA Medical Center resources and facilities to A.A.P.

